# Paternal transmission of behavioural and metabolic traits induced by postnatal stress to the 5^th^ generation in mice

**DOI:** 10.1101/2022.07.11.499529

**Authors:** Chiara Boscardin, Francesca Manuella, Isabelle M Mansuy

**Author notes:** Corresponding author: Isabelle M Mansuy. Laboratory of Neuroepigenetics, University of Zurich and ETH Zurich, Winterthurerstrasse 190, 8057 Zurich, Switzerland.

## Abstract

Life experiences and environmental conditions in childhood can change the physiology and behaviour of exposed individuals and in some cases, of their offspring. In rodent models, stress/trauma, poor diet and endocrine disruptors in a parent have been shown to cause phenotypes in the direct progeny, suggesting intergenerational inheritance. A few models also examined transmission to further offspring and suggested transgenerational inheritance, but such multi-generations inheritance is not well characterized. Our previous work in a mouse model of early postnatal stress showed that behaviour and metabolism are altered in the offspring of exposed males up to the 4^th^ generation in the patriline and up to the 2^nd^ generation in the matriline. The present study examined if in the patriline, symptoms can be transmitted beyond the 4^th^ generation. Analyses of the 5^th^ and 6^th^ generation of mice revealed that altered risk-taking and glucose regulation caused by postnatal stress are still manifested in the 5^th^ generation but are attenuated in the 6^th^ generation. Some of the symptoms are expressed in both males and females, but some are sex-dependent and sometimes opposite. These results indicate that postnatal trauma can affect behaviour and metabolism over many generations, suggesting epigenetic mechanisms of transmission.

## INTRODUCTION

The environment strongly influences physiology in plants and animals. Many environmental factors can modify phenotypes persistently and affect mental and physical health in mammals. They represent serious health risk, and it is estimated that 12.6M global deaths per year are due to modifiable environmental factors [1]. If germ cells are affected, and have alterations that persist until conception, they may be transferred to the embryo at fertilization, and result in symptoms of exposure in the progeny. Such inheritance is not due to changes in the genetic sequence, but likely involve epigenetic factors and mechanisms. Transmission of environmentally-induced traits has been extensively documented in various species including plants, ciliates, chicken, fish, *Neurospora crassa, C. elegans, Drosophila* and mammals [2]–[9]. In humans, large epidemiological studies in historical cohorts such as the Överkalix, Dutch Hunger Winter Famine and ALSPAC have suggested that food supply and famine in prenatal life or childhood of grand-parents have an incidence on the risk to develop cardiovascular diseases, obesity and mortality in descendants [10]–[13]. Likewise, endocrine disruptive chemicals, smoking and led can affect health across generations in humans [14]–[18].

Further to conditions involving food and chemicals, emotional and psychological factors can also strongly affect health. Up to 45% of children in developed countries and over 50% in emerging countries are exposed to adverse experiences such as emotional, physical and sexual abuse, household violence, neglect or parental loss. Such detrimental conditions increase the risk for depression, mood disorders, addiction, and comorbidities such as cardiometabolic and autoimmune diseases, and cancer, in exposed individuals [19]–[21] and their descendants [17], [18], [22], [23]. Childhood trauma is one of the leading causes of premature death in adulthood [24]. While maternal or caregiver behaviour and nursing are possible ways by which the effects of early life adversity can be transferred to children [25]–[27], transmission via gametes is another likely route [28], [29]. The involvement of the parental germline is supported by adoption studies in humans [30] and cross-fostering in animals [31], [32]’and implies that the offspring may be affected independently of parental care.

Several rodent models have been developed to examine the effects of early life stress across generations. We established a model of postnatal trauma in mice based on unpredictable maternal separation combined with unpredictable maternal stress (MSUS) [33]. The model mimics childhood trauma in humans resulting from parental neglect, unreliable care and poor affective attachment. It consists in separating mothers (F0) from their new born pups (F1) each day for 3 hours at an unpredictable time, starting one day after birth until postnatal day (PND) 14 (Fig. 1 A). During separation, females are further exposed unpredictably to a stressor (acute swim or restraint chosen randomly), which further increases distress (Fig. 1 B). When adult, the offspring has behavioural, cognitive and metabolic symptoms that persist across life and are transmitted to the offspring, via both females and males [32]–[39]. Phenotypes include risk-taking behaviours, depressive-like symptoms, altered social abilities, memory deficits, insulin/glucose dysregulation and altered bone homeostasis but also in some conditions, stress resilience and improved behavioural flexibility. Recently, we showed that some of the symptoms of MSUS are transmitted to the 4^th^ generation via males [40].

**Figure 1.**
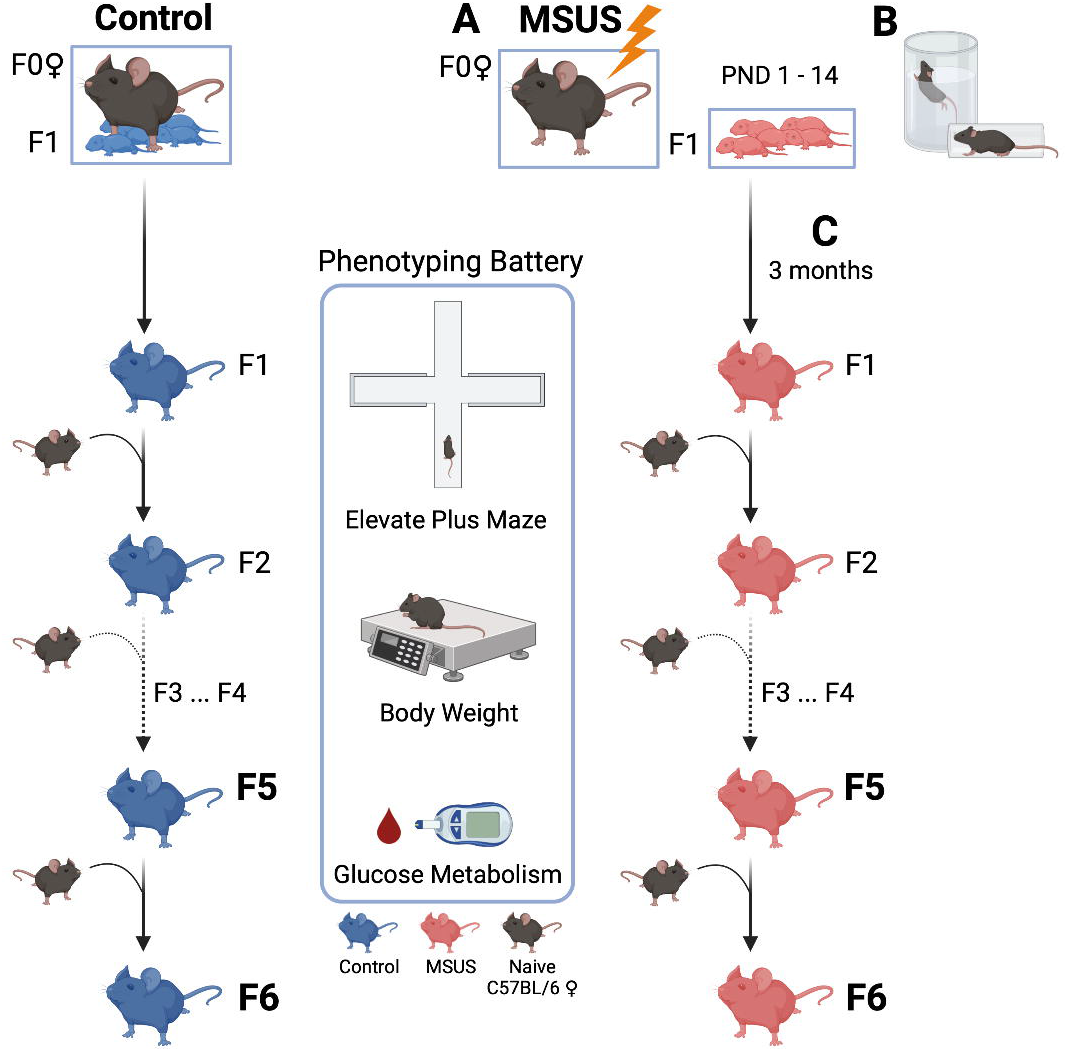
MSUS paradigm. (**A**) For MSUS, naïve males and primiparous females are mated, and starting one day after birth until postnatal day 14 (PND1-14), pups (F1) are separated from their mother (F0) for 3 hours daily at an unpredictable time during the active cycle. (**B**) During separation, dams are exposed to an unpredictable stressor which is either an acute (5-min) swim in 18 °C water or a 20-minute physical restraint in a tube anytime (unpredictably) during the 3 hours. Between PND15 and weaning at PND21, pups are left undisturbed with their mother, and are then raised normally until adulthood. Control animals are produced and raised normally (left). (**C**) When adult (>3 months), F1 males are mated with naïve primiparous control females to produce F2 offspring and breeding is repeated with males of each generation to produce F3, F4, F5 and F6 offspring. Male breeders are removed from the mating cage after one week, and are never in contact with their offspring. MSUS is applied only to F1 pups but not to any of the following offspring. When adult, animals from each generation undergo behavioural and metabolic testing. The present study used F5 and F6 animals. Created with BioRender.com.

To extend our previous findings, we examined if behavioural and metabolic symptoms induced by MSUS can be transmitted to animals beyond the 4^th^ generation. We produced 5^th^ and 6^th^ generation MSUS and control offspring from the patriline by breeding 4^th^ and 5^th^ generation males to naïve females respectively (Fig. 1 C), and assessed their behaviour and metabolic responses. The results show that altered risk-taking behaviours and glucose/insulin responses are still manifested by animals of the 5^th^ generation, in a sex-dependent manner and with opposite effects for glucose/insulin responses. The symptoms are attenuated in animals of the 6^th^ generation.

## MATERIAL AND METHODS

### Mice

C57BI/6JRj mice (Elevage Janvier, Le Genest Saint Isle, France) were maintained in a temperature- and humidity-controlled facility on a 12h reversed light–dark cycle (white light from 8 pm to 8 am, darkness from 8 am to 8 pm), in individually ventilated cages (SealSafe PLUS, Tecniplast, Germany) with food (M/R Haltung Extrudat, Provimi Kliba SA, Switzerland, Cat. #3436) and water *ad libitum*. Cages contained wood chip bedding (LIGNOCEL SELECT, J. Rettenmaier & Söhne), paper tissue as nesting material and a plastic house. All procedures were carried out during the animals’active phase (darkness) in accordance with guidelines and regulations of the cantonal veterinary office in Zurich and the Swiss Animal Welfare Act (Tierschutzgesetz). All animal experiments were approved by cantonal veterinary authorities (license 57/2015 and 83/2018).

### Unpredictable maternal separation combined with unpredictable maternal stress (MSUS)

Naïve males and primiparous C57BI/6JRj females (3-4 months old) were bred in a one to one pairing. After a week, males were removed from the cage to avoid any interference with gestation. At delivery, dams (F0) and pups (F1) were randomly assigned to MSUS or control groups, but taking into consideration the number of male/female pups born in each litter to obtain comparable group size. MSUS pups were separated from their mother daily for 3 hours at an unpredictable time during the active phase starting at postnatal day (PND) 1 until PND14 (Fig. 1 A). During separation, MSUS dams were randomly and unpredictably subjected to either an acute swim in cold water (18°C for 5 minutes) or restraint for 20 minutes in a plastic tube (3.18 cm in diameter with sliding nose restraint and air holes, Midsci) (Fig. 1 B). Control animals were left undisturbed. Both control and MSUS mice had weekly cage changes and weight measurements. Pups were weaned at PND21 and reared in social groups (3–5 mice/cage, controls or MSUS) but from different dams to avoid litter effects. To produce offspring, adult control and MSUS males at each generation were mated with nai□ve primiparous C57BI/6JRj females, and F2, F3, F4, F5 and F6 MSUS offspring were obtained (Fig. 1 C). The MSUS paradigm was applied only to F0 dams and F1 offspring. F2, F3, F4, F5 and F6 MSUS offspring were not exposed to any treatment.

### Breeding size and litter numbers

The number of males used for breeding at each generation depended on the number of offspring needed for the experiments and ranged from 10 to 40 [40]. Litters with 4-10 male and female pups were used for final cage assignment. Litters with less than 4 pups or more than 10 pups were excluded. Only litters used for experiments are reported (Supplementary Table S1). Breeding was conducted after phenotyping, after 2-3 weeks of rest following the last test.

### Behavioural testing

Mice were singly housed 1h before the elevated plus maze to avoid collective stress of cage mates taken away for testing. This reduces experimental variability and improves data quality with small groups of mice [41]. The animals were moved from the colony room to the experimental room immediately before the test. The elevated plus maze consisted of a dark grey PVC platform with two open (length: 30 cm, width: 5 cm) and two closed (with 15 cm walls) arms, elevated 60 cm above the floor. A video camera was placed directly above the centre of the maze and two overhead white lights were positioned to obtain different levels of illumination of open (18±1 Lux) and closed (9±1 Lux) arms. Animals were removed from their home cage by the tail and placed directly into the centre of the maze (5×5 cm), facing the open arm opposite from the experimenter. Tracking/recording lasted 5 minutes and was remotely initiated when the mouse was on the platform. Time spent in open and closed arms, and total distance covered were automatically recorded by a videotracking system (ViewPoint Behaviour Technology). The latency to first enter an open arm was manually scored. Animals that never entered an open arm were assigned a latency of 300 seconds (equal to the duration of the entire task). The maze was cleaned after each animal. The experimenter was blind to the identity and treatment of animals. Testing was conducted in adult animals (3-6-months old) and during the animals’ active phase (darkness). Experiments in males and females were run on different days to avoid confounding effects of olfactory cues between sexes.

### Metabolic assays

Body weight measuraments were conducted under dim red light, while blood sampling was done in white light. Experimenters were blind to the identity and treatment of animals.

#### Food intake and body weight

Food pellets were weighted in each cage every day for 72 hours and averaged by the number of animals (maximum 4 animals/cage). Every 24 hours, pellets were replaced with fresh pellets to limit crumb spillage. Animals were weighed before the experiment and right after the last food measurement. Caloric intake was calculated as the mean amount of food consumed during 72 hours over mean body weight per cage.

#### Glucose level in response to restraint

The test was performed as described in [40]. Each individual mouse was placed for 30 minutes in a cylindrical plastic tube (3.18 cm diameter with sliding nose restraint and air holes; Midsci) for physical restraint and the tail was fixed on the table with tape. Blood was drawn at 0, 15, 30, 90 minutes by tail prick, within 1 cm of tail end using a 22G needle. After 30 minutes, the animal was placed in a temporary cage for one hour. For the 90-minute measurement, each mouse was confined under an inverted 1-litre glass beaker (14 cm high and 12 cm diameter) with its tail protruding from under the spout to allow access by the experimenter for the last blood sampling. The animal was then immediately placed back into its cage. Glucose was measured from fresh blood droplets with an Accu-Chek Aviva glucometer (Roche).

#### Glucose and insulin tolerance test (GTT and ITT)

Mice were temporarily singly housed and fasted for 5 hours before the experiment. Glucose was measured in blood samples at 0, 15, 30, 90 and 120 minutes after an intraperitoneal injection of glucose (GTT) or insulin (ITT). For blood sampling, each animal was confined under an inverted 1 litre glass beaker (14 cm high and 12 cm diameter) with the tail exiting from under the spout. For GTT, 2 mg/g body weight glucose in 0.45% (wt/vol) saline was injected. For ITT, 1 mU of insulin (NovoRapid Novo Nordisk A/S) in sterile 0.9% saline per gram body weight (1mU/g) was administered. If blood glucose fell below 1.7 mM/ml, the animal was rescued with an intraperitoneal injection of 2 mg/gram glucose and removed from the experiment. Since many of the F5 females had to be rescued after the injection of 1mU/g of insulin, a dose of 0.6 mU/g was used for F6 females. For both GTT and ITT, each mouse was placed in a temporary cage after 30 minutes and confined again under a glass beaker 90 minutes after the initial injection to conduct the last two blood samplings. Glucose level was determined in fresh tail blood using an Accu-Chek Aviva glucometer (Roche).

### Statistics

Statistical analyses were performed using Graphpad Prism software, version 8 and 9. Sample size was estimated based on previous experiments on the MSUS model [33], [37], [40]. Data were screened for outliers using Prism’s robust regression and outlier removal test (ROUT, Q set at 5%) for weight, distance covered on the elevated plus maze, and food intake. Animals identified as outliers were excluded from the analysis. For all the other tests, mice were excluded only if technical problems occour (e.g. sudden noise during behavioural testing, interruption of tasks to rescue the animals. Supplementary Table S2). D’Agostino-Pearson and/or Shapiro-Wilk tests were used to assess data distribution. When data followed a Gaussian distribution, parametric tests were used including two-tailed Student’s t-test to compare 2 groups and ANOVAs to compare more than 2 groups. Welch’s correction was applied if the variance was not homogeneous between groups. In the case of non-parametric distribution, Mann-Whitney U test was used to compare 2 groups. Significant effects in ANOVAs were further analyzed using Sidak’s multiple comparison test. Significance was set at p < 0.05 for all tests and indicated by asterisks, and p < 0.1 was considered a trend and indicated by a hashtag. Reported *n* represents data after outliers removal. Control mice are represented in blue, MSUS mice in red.

## RESULTS

### Production of 5th and 6th generation offspring

15 control and 16 MSUS male mice (8.5 month-old) from the 4^th^ generation (F4) [40], obtained by breeding of the grand-offspring (F3) of exposed males (F1), were each paired with one naïve primiparous female to generate F5 offspring (Supplementary Table S1). The breeding produced 14 control and 13 MSUS F5 litters, for a total of 104 males (57 control and 47 MSUS) and 48 females (24 control and 24 MSUS). Likewise, to obtain F6 offspring, 15 control and 16 MSUS F5 males (10 month-old) were each mated one-to-one with a naïve primiparous female and produced 6 control and 10 MSUS litters, with 60 males (30 control and 30 MSUS) and 40 females (20 control and 20 MSUS, Supplementary Table S1).

### Anxious behaviours in MSUS males of the 5th generation

We assessed the response to aversive conditions in F5 and F6 adult animals using an elevated plus maze. F5 MSUS males spent significantly less time in the open arms of the maze than controls (Fig. 2 A) but had a similar latency to first enter an open arm (Fig. 2 B). Notably, the reduced time in open arms is opposite to that previously observed in F1, F2, F3 and F4 males [37], [40]. Time spent in open arms and latency to first enter an open arm in F6 MSUS males were not significantly different from controls despite a trend for an increase (p = 0.0842, Fig. 2 C, D). F5 and F6 MSUS females did not show any behavioural alteration, suggesting no apparent transmission of risk-taking traits beyond F4 (Fig. 2) [40]. Locomotor activity in male and female mice of both F5 and F6 generations was unaffected by MSUS (Supplementary Fig. S1).

**Figure 2.**
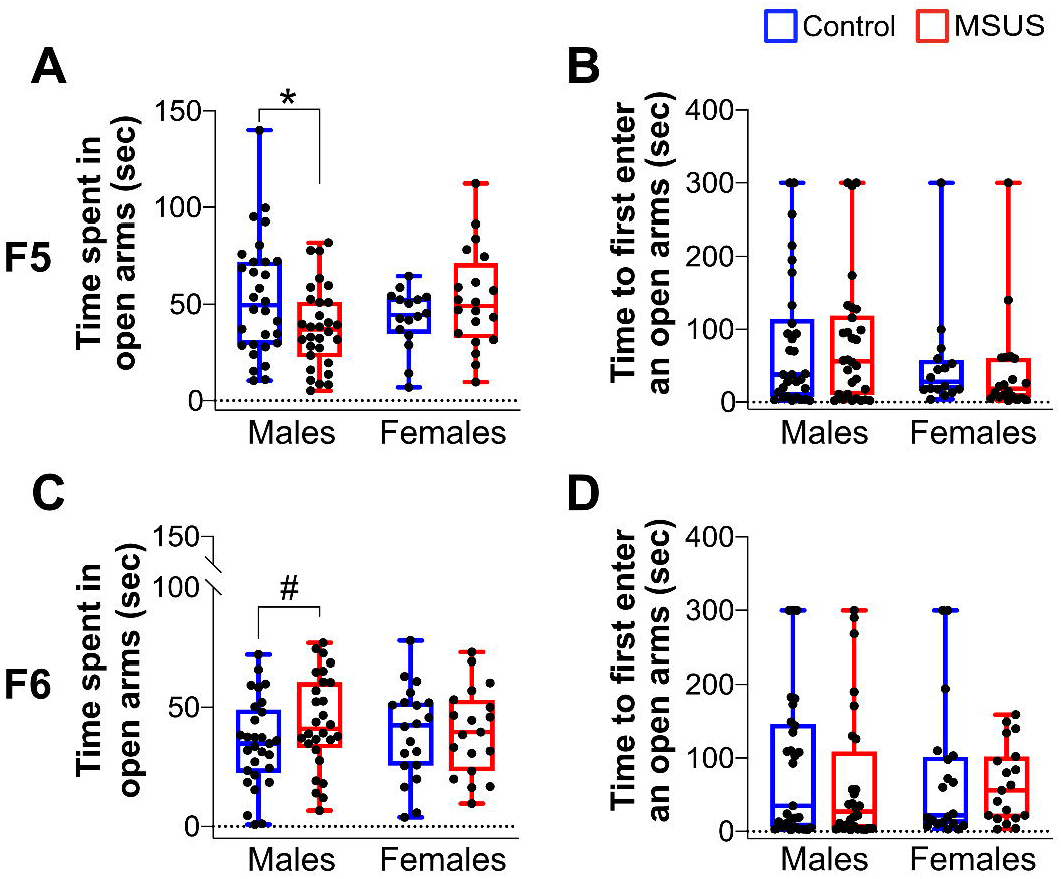
Performance of F5 and F6 adult mice on an elevated plus maze. F5 MSUS males (**A**) spend less time in the open arms of the maze than control males (Controls n = 30, MSUS n = 30, t_58_ = 2.279, p = 0.0263) but (**B**) have comparable latency to first enter an open arm (Controls n = 30, MSUS n = 30, U = 441, p = 0.8978). F5 control and MSUS females (**A**) spend similar time in open arms (Controls n = 16, MSUS n = 20, t_34_ = 1.346, p = 0.1873), and (**B**) have comparable latency to first enter an open arm to control females (Controls n = 16, MSUS n = 20, U = 125, p = 0.2722). (**C, D**) There is a trend for F6 MSUS males to spend more time in open arms than control males (Controls n = 29, MSUS n = 28, t_55_ = 1.759, p = 0.0842) but no difference in females (Controls n = 20, MSUS n = 19, t_37_ = 0.0815, p = 0.9354). Latency to first enter an open arm is comparable in control and MSUS males (Controls n = 29, MSUS n = 28, U = 360, p = 0.4677) and females (Controls n = 20, MSUS n = 19, U = 166.5, p = 0.5177). Data represent median ± whiskers. #p < 0.1, *p < 0.05.

### Altered body weight in MSUS males and females of the 5th generation

F5 MSUS males and females were overweighed from PND7 until early adulthood (4.5 months) when compared with control animals (Fig. 3 A, B). This is consistent with the increased weight observed in adult F4 MSUS males [40] but opposite to the reduced weight in adult MSUS mice from F2 and F3 generations, suggesting a rebound across generations. The overweight phenotype was no longer observed in MSUS females at 7.5 months (Fig. 3 B). We assessed if the increased body weight is due to higher food consumption by measuring caloric intake. F5 MSUS males consumed an amount of food comparable to controls but F5 MSUS females consumed slightly, but not significantly more (p = 0.081) than control females (Fig. 4). This suggested that the increased weight is not due to excessive eating but most likely to a metabolic dysregulation. F6 MSUS animals had normal weight (Fig. 3 C, D) and food intake (data not shown) compared to controls.

**Figure 3.**
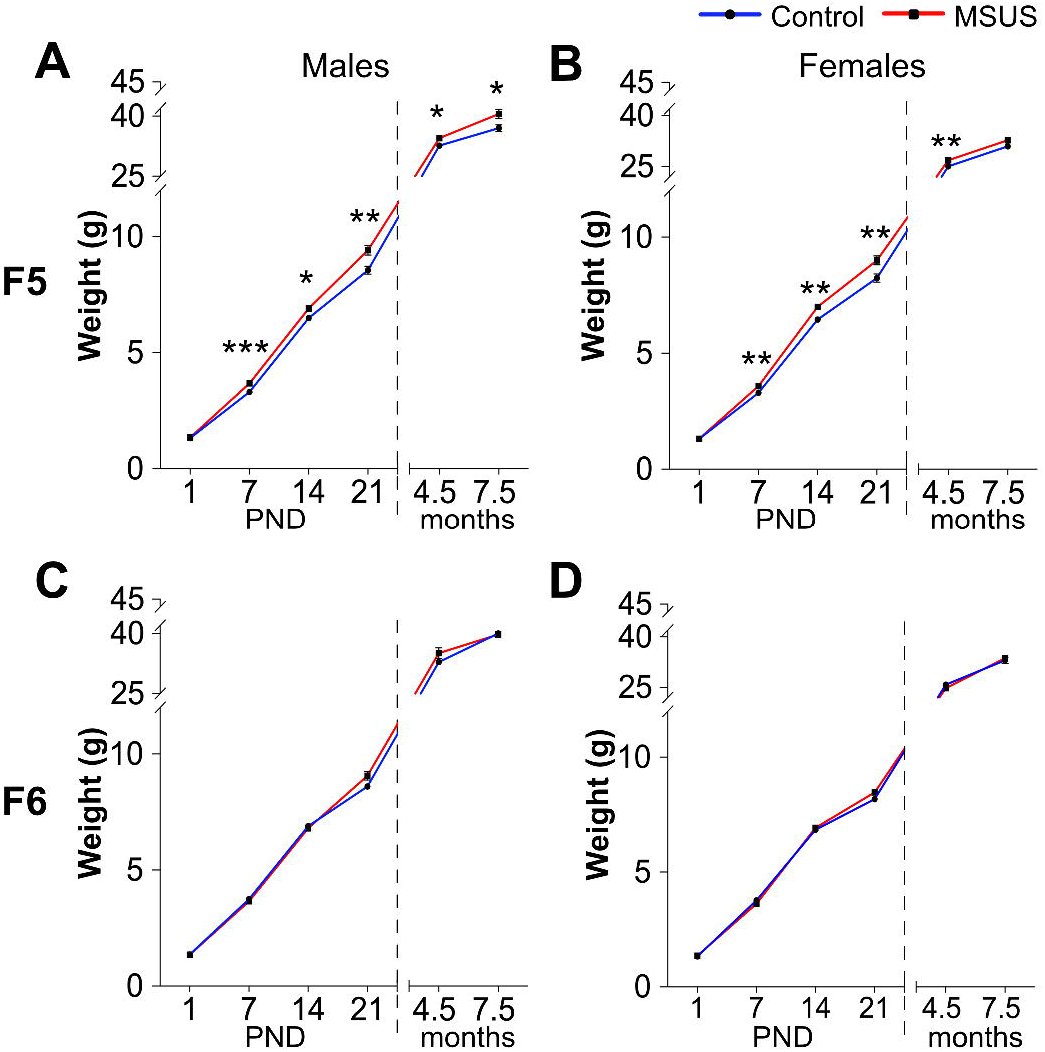
Weight across development in F5 and F6 mice. **(A)** F5 MSUS males weigh more than controls at PND7 (Controls n = 54, MSUS n = 46, t_98_ = 3.932 p = 0.0002), PND14 (Controls n = 56, MSUS n = 46, t_100_ = 2.527, p = 0.0131), PND21 (Controls n = 56, MSUS n = 45, t_99_ = 3.080, p = 0.0027), 4.5 months (Controls n = 28, MSUS n = 22, Welch’s t_28.33_ = 2.331, p = 0.0271) and 7.5 months (Controls n = 16, MSUS n = 16, t_30_ = 2.435, p = 0.0210) but not at PND1 (Controls n = 58, MSUS n = 46, t_102_ = 1.084, p = 0.2808). **(B)** F5 MSUS females weight more than controls at PND7 (Controls n = 41, MSUS n = 42, t_81_ = 3.401, p = 0.0010), PND14 (Controls n = 48, MSUS n = 42, t_88_ = 3.059, p = 0.0029), PND21 (Controls n = 48, MSUS n = 43, t_89_ = 2.887, p = 0.0049) and 4.5 months (Controls n = 17, MSUS n = 20, Welch’s t_26.44_ = 3.068, p = 0.0049) but not at PND1 (Controls n = 49, MSUS n =46, t_93_ = 0.0074, p = 0.9940) or at 7.5 months (Controls n = 18, MSUS n = 20, t_36_ = 1.614, p = 0.1153). No weight differences are observed in F6 MSUS (**C**) males (PND1 controls n = 35, MSUS n = 50, t_83_ = 0.0716, p = 0.9431; PND7 controls n = 34, MSUS n = 51, U = 806.5 p = 0.5907; PND14 controls n = 34, MSUS n = 42, t_74_ = 0.3191, p = 0.7506; PND21 controls n = 34, MSUS n = 42, t_74_ = 1.164, p = 0.2481; 4.5 months controls n = 16, MSUS n = 16, t_30_ = 1.392, p = 0.1741; 7.5 months controls n = 16, MSUS n = 16, t_30_ = 0.0015, p = 0.9988) or (**D**) females (PND1 controls n = 25, MSUS n = 69, t_92_ = 1.042, p = 0.3003; PND7 controls n = 29, MSUS n = 66, U = 894, p = 0.6140; PND14 controls n = 24, MSUS n = 54, U = 570, p = 0.4025; PND21 controls n = 23, MSUS n = 53, Welch’s t_56.75_ = 1.663, p = 0.1011; 4.5 months controls n = 15, MSUS n = 15, t_28_ = 1.692, p = 0.1018; 7.5 months controls n = 20, MSUS n = 20, t_38_ = 0.5016, p = 0.6189). Numbers variation among PND1-7-14-21 timepoints might be due to outliers’ removal (see also Material and Methods, Statistics), wrong gender identification, perinatal death. For adult mice (4.5 and 7.5 months), only a subset of the total number of animals underwent weight measurements. Data represent mean ± s.e.m. #p < 0.1, *p < 0.05, **p < 0.05, ***p <0.001.

**Figure 4.**
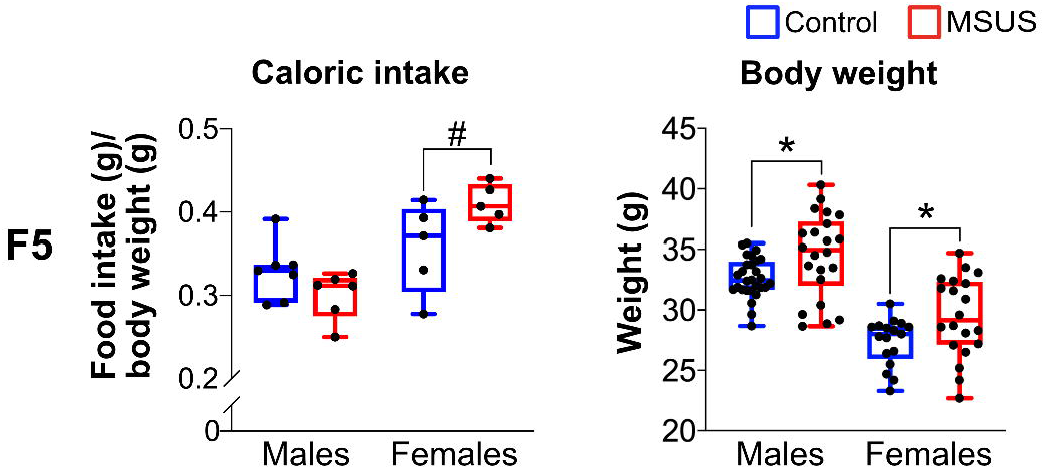
Caloric intake in adult F5 mice. The average caloric intake/cage is comparable in F5 control and MSUS males (Controls n = 7 cages, MSUS n = 6 cages, t_11_ = 1.578, p = 0.1429, 4.5 months) and females (Controls n = 5 cages, MSUS n = 5 cages, t_8_ = 1.996, p = 0.0810, 6 months) but weight is higher in F5 MSUS than control males (Controls n = 28, MSUS n = 22, Welch’s t_28.33_ = 2.331, p = 0.0271) and females (Controls n = 17, MSUS n = 20, Welch’s t_31.45_ = 2.347, p = 0.0254). Data represent median ± whiskers. #p < 0.1, *p < 0.05.

### Dysregulated glucose homeostasis in F5 and F6 MSUS mice

We examined glucose response in F5 and F6 animals using a glucose tolerance test (GTT) [42]. Glucose level in F5 MSUS females but not males was lower than controls at the peak of the curve, suggesting that MSUS females have a more efficient glucose clearance (Fig. 5 A, B and Supplementary Table S3). We then used an insulin tolerance test (ITT) to assess glucose response after an insulin injection [42]. Insulin provoked a more pronounced decline in blood glucose in F5 control than MSUS males, even if the overall trend of the response was not changed (Fig. 5 C, Supplementary Table S3). F5 females did not show any difference between control and MSUS in the ITT (Fig. 5 D, Supplementary Table S3) although only half of the animals completed the test (6 controls, 7 MSUS) and the other half had to be rescued by glucose injection due to signs of distress. Consistent with F5 MSUS males, F6 MSUS males had a lower glucose response to insulin during ITT (Fig. 5 E, Supplementary Table S4) but no change during GTT (Supplementary Table S3). No alteration in glucose response after GTT or ITT was observed in F6 MSUS females (Fig. 5 F, Supplementary Table S4). We next examined glucose response after an acute stress using a restraint test. Although blood glucose was transiently increased in F5 and F6 MSUS and control animals, there was no significant overall difference between the groups (Supplementary Table S3 and S4). This suggests that the blunted increase in blood glucose observed in F2 and F4 MSUS mice [37], [40] is not transferred to subsequent generations or is compensated for.

**Figure 5.**
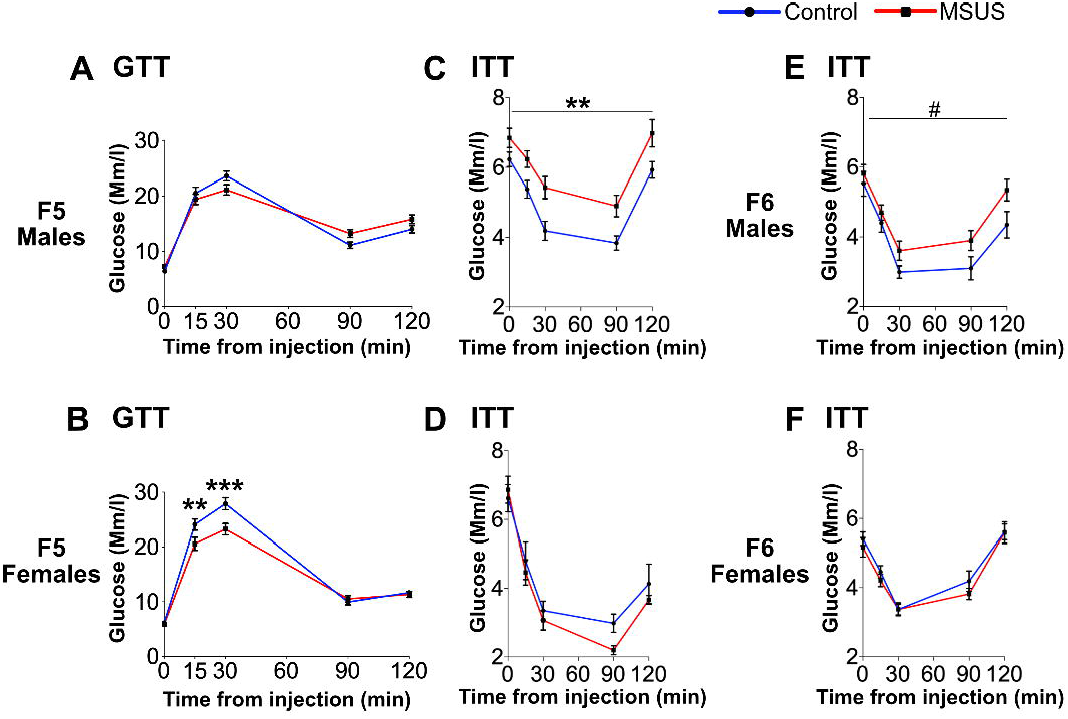
Metabolic assessment in F5 and F6 mice. **(A, B)** In the glucose tolerance test, **(A)** F5 control and MSUS males have comparable level of blood glucose (Controls n = 15, MSUS = 14, interaction F(4,108) = 4.469, p = 0.0022; post-hoc test: p > 0.05) but **(B)** F5 MSUS females have lower blood glucose at the peak of the curve than controls (Controls n = 13, MSUS = 14, interaction F(4,100) = 4.423, p = 0.0025; post-hoc test: p_T15_ = 0.0098; p_T30_ = 0.0003). (**C-F**) In the insulin tolerance test, insulin response is **(C)** significantly blunted in F5 MSUS males compared to controls (Controls n = 16, MSUS = 15, group effect: F(1, 29) = 10.17, p = 0.0034) but **(E)** not in F6 MSUS males (Controls n = 12, MSUS = 14, group effect F(1,24) = 3.549, p = 0.0718) or (**D**) in F5 (Controls n = 6, MSUS = 7) and (**F**) F6 (Controls n = 13, MSUS = 14) females. Data represent mean ± s.e.m. #p < 0.1, *p < 0.05, **p < 0.05, ***p < 0.001. Detailed results are presented in Supplementary Table S3 (F5) and S4 (F6).

## DISCUSSION

The current study extends our previous findings that postnatal traumatic stress in mice causes behavioural and metabolic changes that persist across 3 generations of offspring (till F4). The MSUS model is characterized by increased risk-taking, which is the most penetrant trait observed in directly exposed males and their offspring up to the 4^th^ generation [40]. This trait was however not manifested by animals of the 5^th^ generation but instead, F5 males showed increased anxiety. Mice of the 6^th^ generation had nonetheless a trend for increased risk-taking. Notably, we observed in the past that some MSUS traits are not manifested in one generation but are observed in the offspring e.g. depressive-like behaviour not detected in F2 but in F3 mice [33]. The advanced age of F5 fathers at breeding (10 months) might also contribute to the attenuation of the MSUS phenotype from the 5^th^ to the 6^th^ generation, and to the reduced number of F6 litters as well.

Animals of the 5^th^ generation have an altered glucose homeostasis similarly to mice from previous generations [37], [40]. The reduced glucose level observed on GTT in MSUS females may involve different mechanisms including upregulated glucose effectiveness in stimulating its own uptake and suppressing its own production under basal/constant insulin concentrations [42]–[44]. The increased glucose level on ITT in MSUS males may also result from a reduced sensitivity to insulin or insulin resistance, suggesting a pathological state similar to metabolic syndrome [45] or type 2 diabetes [46]. It may also be due to changes in other factors such as insulin counter-regulatory hormones like glucagon, epinephrine, cortisol or growth hormone, which sustain plasma glucose during fasting conditions [47]. Additional methods such as glucose clamp technique or a direct quantification of insulin level would be required to fully assess changes in glucose metabolism in MSUS animals. Our results show that glucose response is not altered in animals of the 6^th^ generation, suggesting that the underlying factors of transmission in the germline were compensated for or corrected, perhaps progressively across generations. The fact that MSUS males are bred with non-exposed naïve females at each generation probably favours such correction. The manifestation of metabolic phenotypes after early life trauma that persist across generations is highly relevant to humans, and is reminiscent of the increased risk for physical health problems in people exposed to childhood trauma [48]. Other models of exposure e.g. to endocrine disruptors were reported to have phenotypes across generations [49].

The mechanisms and evolutionary benefits of such transmission are not fully understood and may be different depending on the type of exposure. Challenging environmental conditions can increase fitness by inducing phenotypic adaptation to these conditions. But, if the conditions change, the progeny is then mal-adapted and may express deficits or negative phenotypes [50]. Further, not all phenotypes have the same depth of penetrance, causing some features to be transmitted from exposed parent to direct offspring but not further. The scarcity of mammalian models investigating the transgenerational effects of environmental exposure beyond the 2^nd^ or 3^rd^ generation limits the understanding of these effects. Among these models, which include prenatal stress [51], environmental toxicants [2], obesogens [52], drugs [53], [54], the MSUS model presents several advantages and unique features. It allows study both inter-and transgenerational inheritance, and relies on a non-invasive manipulation, highly similar to aspects of childhood mistreatment in humans. Unlike many models which have extensive exposure spanning preconception to postnatal or even adult life [55]–[58], MSUS is based on a short and time-restricted treatment in postnatal life (PND1 to PND14). This treatment does not interfere with prenatal development or epigenetic reprogramming during embryogenesis [29]. The restricted time window makes it easier to know which cells are present during these developmental stages and are likely to be affected by exposure. For instance, developing gonads in mice mostly have spermatogonial cells during early postnatal life [29], [59], that may be directly affected. Further, the patriline design helps avoid confounding factors such as the intrauterine milieu and maternal behaviours that can mask true germline-dependent transmission [41], [60]– [62]. It allowed us to identify sperm as a carrier of molecular signals of exposure from father to offspring [37]. So far, sperm RNA is one the best documented vectors of epigenetic inheritance [7], that we demonstrated to be causally responsible for the transmission of the effects of MSUS to the offspring [37], [63]. Recently, we also provided evidence that some of the effects of exposure in the offspring can be recapitulated by intravenous injection of MSUS serum in adult control males [64], suggesting that blood components can influence the germline [4], [65], [66]. Developing other models of exposure with both patriline and matriline transmission would help better understand germline-dependent mechanisms of epigenetic inheritance.

## Authors’ contribution

CB, FM and IMM conceived and designed the study. CB and FM conducted MSUS paradigm, performed behavioural and metabolic experiments and collected the results. FM organized and managed breeding. CB compiled and analysed raw data, and prepared figures. CB, FM and IMM wrote the manuscript. IMM raised funds to support this project.

## Acknowledgments

We thank the University Zürich, the Swiss Federal Institute of Technology Zürich, the Swiss National Science Foundation (grant number 31003A_175742/1), EU Horizon 2020 Research and Innovation Program Grant N° 848158 EarlyCause, Escher Family Fund for supporting this research. We thank Gretchen van Steenwyk and Rodrigo Arzate for critical discussion of the results, Alberto Corcoba and Deepak Tanwar for help with statistical analyses, and Yvonne Zipfel and the LASC team for excellent animal care.

## Data availability

The data that support the findings of this study are available from the corresponding author, IMM, upon reasonable request.

