## Supplementary for "Paternal transmission of behavioural and metabolic traits induced by postnatal stress to the 5^th^ generation in mice"

Supplementary Figure S1 (Boscardin et al.)

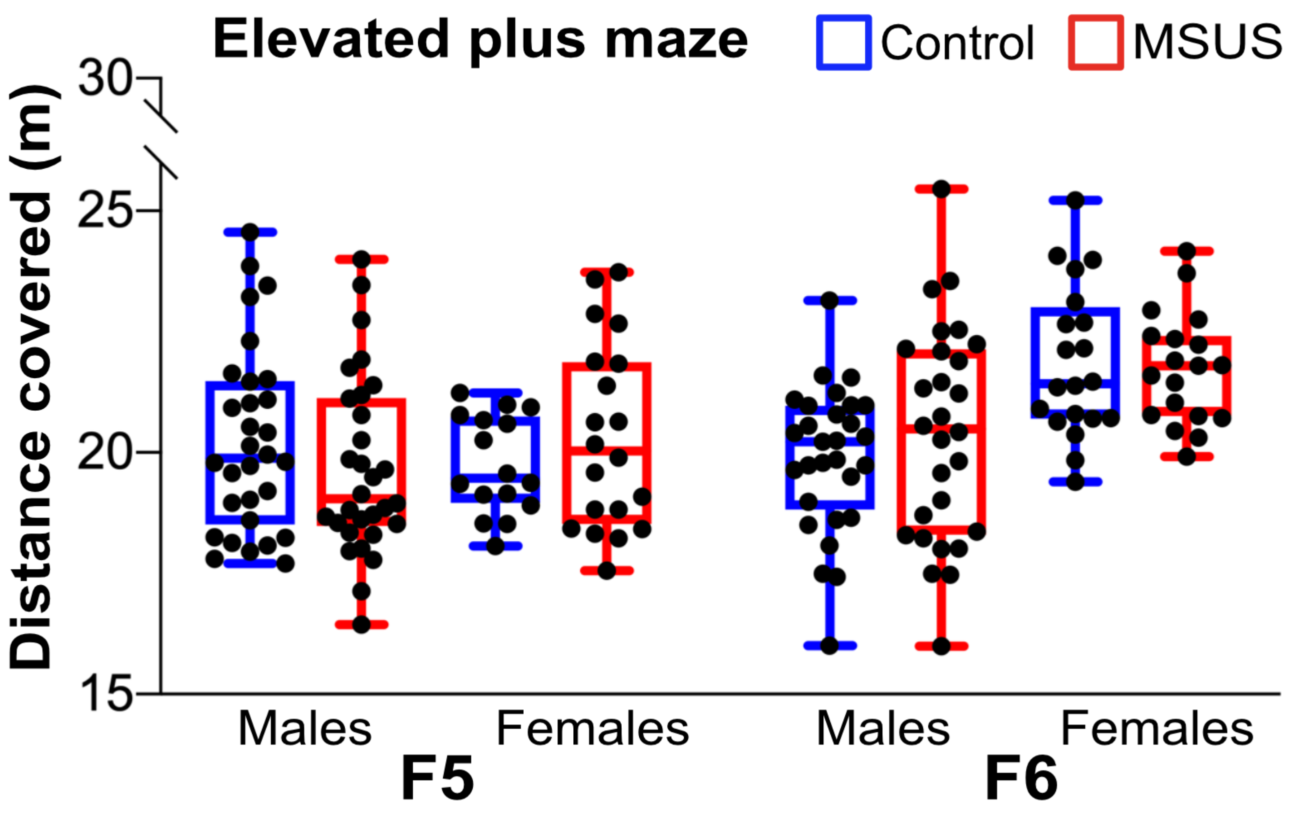


**Supplementary Figure S1. Locomotor activity on the elevated plus maze (Figure 2).** F5 and F6 control and MSUS mice show a comparable locomotor activity on the elevated plus maze. F5 males: Controls n = 30, MSUS n = 30, $t_{58}$ = 1.152, p = 0.2540; F5 females: Controls n = 16, MSUS n = 20, Welch's $t_{30}$ = 1.149, p = 0.2598. F6 males: Controls n = 29, MSUS n = 28, Welch's $t_{46.76}$ = 0.9882, p = 0.3281; F6 females: Controls n = 20, MSUS n = 19, $t_{37}$ = 0.2667, p = 0.7912. Data represent median ± whiskers.

Supplementary Table S1 (Boscardin et al.)

| | Breeding F4 → F5 | | Breeding F5 → F6 | |
|---|---|---|---|---|
| | Control | MSUS | Control | MSUS |
| **Breeding pairs** | 15 | 16 | 15 | 16 |
| **Litters at weaning** | 14 | 13 | 6 | 10 |
| **Excluded litters (n < 4 pups)** | - | 1 | 1 | - |
| **Pups at weaning** | 81 | 71 | 50 | 50 |
| **Breeding sucesss rate*** | 90% | | 80% | |
| **Sire age at breeding** | 8.5 months | | 10 months | |
| **Experimenters** | CB, FM | | CB, FM | |

*Number of females that successfully gave birth

**Supplementary Table S1. Technical parameters of MSUS breedings**. Table represents numbers of Control and MSUS breeders used to generate the F5 and F6 generations, information about litters and experimenters involved in the managing of the breedings.

Supplementary Table S2 (Boscardin et al.)

**F5**

| | | MALES | | FEMALES | |
|---|---|---|---|---|---|
| | | Number of mice tested | **Number of mice after experimental outliers exclusion** | Number of mice tested | **Number of mice after experimental outliers exclusion** |
| **EPM** | Control | 31 | **30** | 18 | **16** |
| | MSUS | 31 | **30** | 20 | **20** |
| **GTT** | Control | 16 | **15** | 15 | **13** |
| | MSUS | 16 | **14** | 16 | **14** |
| **ITT** | Control | 16 | **16** | 16 | **6** |
| | MSUS | 16 | **15** | 16 | **7** |
| **Restraint stress** | Control | 16 | **16** | 15 | **15** |
| | MSUS | 16 | **16** | 16 | **16** |

**F6**

| | | MALES | | FEMALES | |
|---|---|---|---|---|---|
| | | Number of mice tested | **Number of mice after experimental outliers exclusion** | Number of mice tested | **Number of mice after experimental outliers exclusion** |
| **EPM** | Control | 30 | **29** | 20 | **20** |
| | MSUS | 30 | **28** | 20 | **19** |
| **GTT** | Control | 16 | **12** | 15 | **12** |
| | MSUS | 16 | **13** | 15 | **14** |
| **ITT** | Control | 16 | **12** | 15 | **13** |
| | MSUS | 16 | **14** | 15 | **14** |
| **Restraint stress** | Control | 16 | **16** | 15 | **15** |
| | MSUS | 16 | **16** | 15 | **15** |

**Supplementary Table S2. Exclusion of experimental outliers in the elevated plus maze, glucose and insulin tolerance test, and restraint stress test.** Table indicates number of animals tested and total number of animals used for statistical analysis, after outliers' exclusion (See Material and Methods, Statistics).

Supplementary Table S3 (Boscardin et al.)

| | F5 males | | | | |
|---|---|---|---|---|---|
| | Number of mice | Source of variation | F (DFn, DFd) | P value | Summary |
| GTT | Control = 15<br>MSUS = 14 | Interaction | F (4, 108) = 4.469 | P=0.0022 | ** |
| | | Time points | F (4, 108) = 177.5 | P<0.0001 | **** |
| | | Group | F (1, 27) = 0.1173 | P=0.7347 | ns |
| ITT | Control = 16<br>MSUS = 15 | Interaction | F (4, 116) = 0.7646 | P=0.5504 | ns |
| | | Time points | F (4, 116) = 51.93 | P<0.0001 | **** |
| | | Group | F (1, 29) = 10.17 | P=0.0034 | ** |
| Restraint stress | Control = 16<br>MSUS = 16 | Interaction | F (3, 90) = 0.4745 | P=0.7008 | ns |
| | | Time points | F (3, 90) = 93.72 | P<0.0001 | **** |
| | | Group | F (1, 30) = 0.04613 | P=0.8314 | ns |

| | F5 females | | | | |
|---|---|---|---|---|---|
| | Number of mice | Source of variation | F (DFn, DFd) | P value | Summary |
| GTT | Control = 13<br>MSUS = 14 | Interaction | F (4, 100) = 4.423 | P=0.0025 | ** |
| | | Time points | F (4, 100) = 236.8 | P<0.0001 | **** |
| | | Group | F (1, 25) = 9.666 | P=0.0046 | ** |
| ITT | Control = 6<br>MSUS = 7 | Interaction | F (4, 44) = 0.4898 | P=0.7432 | ns |
| | | Time points | F (4, 44) = 40.37 | P<0.0001 | **** |
| | | Group | F (1, 11) = 0.01956 | P=0.8913 | ns |
| Restraint stress | Control = 15<br>MSUS = 16 | Interaction | F (3, 87) = 0.6697 | P=0.5729 | ns |
| | | Time points | F (3, 87) = 93.70 | P<0.0001 | **** |
| | | Group | F (1, 29) = 0.5710 | P=0.4560 | ns |

**Supplementary Table S3. Statistics for glucose and insulin tolerance test, and restraint stress in F5**. Table provides detailed results from the metabolic assessment analyses of F5 mice (Figure 5).

Supplementary Table S4 (Boscardin et al.)

| | F6 males | | | | |
|---|---|---|---|---|---|
| | **Number of mice** | **Source of variation** | **F (DFn, DFd)** | **P value** | **Summary** |
| GTT | Control = 12<br>MSUS = 13 | Interaction | F (4, 92) = 0.2294 | P=0.9212 | ns |
| | | Time points | F (4, 92) = 309.5 | P<0.0001 | **** |
| | | Group | F (1, 23) = 0.2147 | P=0.6475 | ns |
| ITT | Control = 12<br>MSUS = 14 | Interaction | F (4, 96) = 1.118 | P=0.3528 | ns |
| | | Time points | F (4, 96) = 45.17 | P<0.0001 | **** |
| | | Group | F (1, 24) = 3.549 | P=0.0718 | # |
| Restraint stress | Control = 16<br>MSUS = 16 | Interaction | F (3, 90) = 0.9356 | P=0.4269 | ns |
| | | Time points | F (3, 90) = 66.18 | P<0.0001 | **** |
| | | Group | F (1, 30) = 0.1966 | P=0.6607 | ns |

| | F6 females | | | | |
|---|---|---|---|---|---|
| | **Number of mice** | **Source of variation** | **F (DFn, DFd)** | **P value** | **Summary** |
| GTT | Control = 12<br>MSUS = 14 | Interaction | F (4, 96) = 0.5717 | P=0.6838 | ns |
| | | Time points | F (4, 96) = 344.3 | P<0.0001 | **** |
| | | Group | F (1, 24) = 0.5295 | P=0.4739 | ns |
| ITT | Control = 13<br>MSUS = 14 | Interaction | F (4, 100) = 0.5385 | P=0.7078 | ns |
| | | Time points | F (4, 100) = 47.84 | P<0.0001 | **** |
| | | Group | F (1, 25) = 0.8258 | P=0.3722 | ns |
| Restraint stress | Control = 15<br>MSUS = 15 | Interaction | F (3, 84) = 0.9257 | P=0.4321 | ns |
| | | Time points | F (3, 84) = 77.65 | P<0.0001 | **** |
| | | Group | F (1, 28) = 1.101 | P=0.3031 | ns |

**Supplementary Table S4. Statistics for glucose and insulin tolerance test, and restraint stress in F6.** Table provides detailed results from the metabolic assessment analyses of F6 mice (Figure 5).
